# Light and growth form interact to shape stomatal ratio among British angiosperms

**DOI:** 10.1101/163873

**Authors:** Christopher D. Muir

**Affiliations:** Biodiversity Research Centre and Botany Department, University of British Columbia, Vancouver, British Columbia V6T 1Z4, Canada

**Keywords:** Adaptation, amphistomy, Ellenberg light indicator value, growth form, phylogenetic comparative methods, Raunkiær life form, stomata, stomatal ratio

## Abstract

- In most plants, stomata are located only on the abaxial leaf surface (hypostomy), but many plants have stomata on both surfaces (amphistomy). High light and herbaceous growth form have been hypothesized to favor amphistomy, but these hypotheses have not been rigourously tested together using phylogenetic comparative methods.
- I leveraged a large dataset including stomatal ratio, Ellenberg light indicator value, growth form, and phylogenetic relationships for 372 species of British angiosperms. I used phylogenetic comparative methods to test how light and/or growth form influence stomatal ratio and density.
- High light and herbaceous growth form are correlated with amphistomy, as predicted, but they also interact; the effect of light is pronounced in therophytes (annuals) and perennial herbs, but muted in phanerophytes (shrubs and trees). Furthermore, amphistomy and stomatal density evolve together in response to light.
- Comparative analyses of British angiosperms reveal two major insights into physiological evolution. First, light and growth form interact to shape variation in stomatal ratio; amphistomy is common under high light, but mostly for herbs. Second, coordinated evolution of adaxial stomatal density and light tolerance indicates that amphistomy is an important adaptation to optimally balance light acquisition with gas exchange. These results advance our understanding of why stomatal ratio evolves and its potential as a functional trait for paleoecology and crop improvement.

## Introduction

Natural selection shapes leaf anatomy in order to optimize its photosynthetic function in a given environment (Haberlandt, 1914; Givnish, 1987; Smith et al., 1997). By understanding the adaptive significance of leaf anatomical variation we can learn about natural history, find targets for crop improvement, and identify anatomical proxies for paleoclimates preserved in the fossil record (e.g. Wolfe, 1971; Royer, 2001; McElwain and Steinthorsdottir, 2017). The size, density, and distribution of stomata on a leaf vary widely and impact the flux of CO_2_ and water vapour (recently reviewed in Sack and Buckley, 2016), as well as susceptibility to foliar pathogens that infect through stomata (McKown et al., 2014; Melotto et al., 2017). Hence, stomata have been especially useful in understanding plastic and evolutionary response to climate change and domestication (Woodward, 1987; Beerling and Royer, 2011; Milla et al., 2013).

While the density and size of stomata have been researched extensively (Sack and Buckley, 2016, and references therein), the adaptive significance of stomatal distribution is less well understood. Stomata are most often found only on the lower leaf surface (hypostomy) but occur on both surfaces (amphistomy) in many species (Metcalfe and Chalk, 1950; Parkhurst, 1978; Mott et al., 1984). Theory and experiments demonstrate that amphistomy increases photosynthetic rates under many conditions. By creating a second parallel pathway for CO_2_ diffusion within the mesophyll, amphistomy optimally supplies CO_2_ (Parkhurst, 1978; Gutschick, 1984; Jones, 1985). Amphistomy is correlated with greater CO_2_ diffusion (Beerling and Kelly, 1996) and higher photosynthetic rates (McKown et al., 2014). These observations are corroborated by experiments demonstrating that amphistomy increases maximum photosynthetic rates by up to 20% (Parkhurst and Mott, 1990). On the other hand, amphistomy can increase transpiration (Jones, 1985; Foster and Smith, 1986; Buckley et al., 2015). While transition to amphistomy is thus thought to increase transpiration, empirical studies suggest greater water-use efficiency in amphistomatous species (Bucher et al., 2017). Hence, amphistomy appears to benefit a plant’s carbon use relative to water loss and should be favored when CO_2_ limits photosynthetic rate. The open questions are under what ecological conditions does CO_2_ supply most strongly limit photosynthetic rate (Peat and Fitter, 1994) and when is photosynthetic rate most important to fitness?

The leading, nonmutually exclusive hypotheses are that 1) open habitats favour amphistomy because CO_2_ diffusion most strongly limits photosynthetic rate under high light and 2) herbaceous growth form favours amphistomy because traits that maximize photosynthetic rate are often under stronger selection in herbs. Salisbury (1927) first noted that amphistomy is most common in herbaceous plants from open habitats (i.e., with high light) of the British flora. These observations have been replicated in other studies (Mott et al., 1984; Peat and Fitter, 1994; Jordan et al., 2014; Muir, 2015) and may support physiological and ecological hypotheses that CO_2_ most strongly limits photosynthesis in high light and/or photosynthesis contributes most to fitness in herbaceous plants. Under high light, CO_2_ can strongly limit maximum photosynthetic rates, especially in thick leaves (Jones, 1985). Hence, having stomata on both surfaces relieves this limitation by adding a second parallel pathway for CO_2_ diffusion. Parkhurst (1978) argued that greater leaf thickness *per se* selected for amphistomy, but there is little evidence for correlations between leaf thickness and stomatal ratio independent of light (Mott et al., 1984; Gibson, 1996; Muir, 2015). Amphistomy is correlated with open habitat in warm desert plants of western North America (Mott et al., 1984; Gibson, 1996), among the Proteaceae (Jordan et al., 2014), and in continental European herbs (Bucher et al., 2017).

Stomatal ratio is also associated with growth form. In the British flora, Salisbury (1927) found that trees and shrubs are nearly always hypostomatous, whereas herbs from open habitats are amphistomatous. This pattern holds when data are averaged by family to coarsely control for phylogenetic nonindependence (Peat and Fitter, 1994) or when using alternative classification schemes, such as Raunkiær life form (Peat and Fitter, 1994). Across plants from ~ 90 families worldwide, growth form is the strongest predictor of stomatal ratio when multiple factors are estimated simultaneously and controlling for phylogenetic nonindependence (Muir, 2015). These patterns are consistent with other data indicating that many herbaceous plants are under strong selection for high maximum photosynthetic rates (Bazzaz, 1979; Körner et al., 1989; Wullschleger, 1993).

Although previous comparative studies have tested whether open habitat and growth form influence stomatal ratio, we do not know if these effects are independent of one another. Open habitat and growth form may be confounded because open habitats generally consist of more short-statured, herbaceous plants. Some authors have attempted to disentangle light and growth form by contrasting herbs from open and understory habitats (Salisbury, 1927). However, this is problematic if phylogenetic relationships are not controlled for, because shade species may share traits simply because they are more closely related to each other than they are to high light species. Finally, open habitat and growth form may also interact with one another. For example, amphistomy may only be favored when CO_2_ strongly limits photosynthetic rate (e.g. in high light) *and* photosynthetic rate strongly limits fitness (e.g. in herbs).

To better understand the adaptive significance of stomatal ratio, I asked three main questions:

1. Are light habitat and growth form correlated?
2. Do light habitat and growth form influence stomatal ratio additively, or do their effects interact?
3. Is evolution of stomatal ratio mediated by changes in stomatal density on the adaxial (upper) surface, abaxial (lower) surface, or both?

In answering these questions, I both reassessed previous hypotheses using newer phylogenetic comparative methods and evaluated previously untested hypotheses. I predicted *a priori* that light habitat and growth would be correlated. Species with faster life histories, especially therophytes (annuals), would on average inhabitat sunnier environments than species with slower life histories, especially phanerophytes (shrubs and trees). Based on hypotheses from previous studies, I also predicted that herbaceous growth form and high light would be associated with amphistomy, even after controlling for phylogenetic nonindependence. Although these predictions have been tested previously, it is critical to reevaluate them here with updated methods because the subsequent untested hypotheses build on these results. The first novel hypothesis I tested predicts that light and growth form interact. Specifically, I hypothesized that both high light and herbaceous growth would be required to favor a more even stomatal ratio (i.e. amphistomy). Finally, I tested whether amphistomy is part of a coordinated syndrome of traits that promote higher photosynthetic rate. If high light and growth form favor amphistomy because it increases photosynthesis, then it follows that they should also favor other stomatal traits that reinforce this advantage. If evolved increases in stomatal ratio are mediated by shifting abaxial stomata to the adaxial surface, holding total stomatal density constant, then the overall increase in CO_2_ diffusion would be small. In contrast, if amphistomy evolves by increasing adaxial stomatal density while holding abaxial density constant, then *total* stomatal density must increase as well. Evolutionary coordination of amphistomy and high stomatal density would thus reinforce one another, increasing CO_2_ supply to chloroplasts more than changes in either trait would in isolation. Understanding selection on coordinated traits can explain the evolution of major functional trait axes and syndromes.

To address these questions, I reanalyzed existing data on stomatal ratio, light habitat, and growth form in British angiosperms (Salisbury, 1927; Fitter and Peat, 1994, 2017) using phylogenetic comparative methods. The British angiosperm flora is well suited for these questions because this flora has been comprehensively surveyed for many ecologically important traits, meaning it is probably the least biased survey of stomatal trait variation. Salisbury’s observations on stomata and ecology in the British flora have heavily influenced plant ecophysiology, but many of his and subsequent authors’ analyses have significant limitations because of inadequate statistical methods. For example, few analyses until recently account for phylogenetic nonindependence (Felsenstein, 1985), which can strongly influence inferences on stomatal traits and growth form (Kelly and Beerling, 1995, this study did not consider light). A species-level phylogeny of the entire British flora (Lim et al., 2014) now allows for the first time a rigorous analysis of evolutionary relationships among stomatal ratio, light, and growth form.

## Materials and Methods

Data and annotated source code to generate this manuscript are available on GitHub (https://github.com/cdmuir/britstom) and Dryad (Muir, 2017).

### Data on stomatal ratio, light habitat, growth form, and phylogenetic relationships

I obtained data on ab- and adaxial stomatal density on 395 species from British Ecological Flora (Salisbury, 1927; Fitter and Peat, 1994, 2017). Following recent comparative analyses (e.g. Niinemets and Valladares, 2006; Bartelheimer and Poschlod, 2016; Shipley et al., 2017), I used Ellenberg light indicator values (Ellenberg, 1974) as measures of light habitat. Hence, I am assuming that the species’ light habitat is closely related to the type of habitat (open versus closed) where that species is found. Ellenberg light indicator values, hereafter abbreviated L-value, have been recently updated by taxonomic experts of the British flora (PLANTATT, Hill et al. (2004)).

There is no universally adopted scientific classification scheme for plant growth form, therefore I statistically competed two widely used schemes based on plant habit and Raunkiær life form. First, I used PLANTATT data on perennation, woodiness, and height to classify species’ growth form based on habit. I categorized herbaceous species as annual, biennial, or perennial and woody species as shrub or tree. Following Muir (2015), ‘biennial’ includes true biennials as well as species that have a mix of perennation forms (e.g. a species with both annual and perennial forms would be classified as a biennial here). Woody species are shrubs (plant height less than 4 m) or trees (plant height greater than 4 m). Next, I compared this scheme to PLANTATT data on Raunkiær life form (Raunkiær, 1934), which is another way to classify growth form in comparative ecology (e.g. Peat and Fitter, 1994; Salguero-Gómez et al., 2016). I retained phanerophytes, geophytes, chamaephytes, hemicryptophytes, and therophytes, but excluded data on hyrdrophytes (14 species) because many of these species are hyperstomatous (Fig. S1) due to the fact that leaves may rest on the water’s surface, selecting for stomata to be present on the upper surface only. The two main differences between these growth form classifications are that 1) most shrubs and trees are lumped together as phanerophytes and 2) many geophytes and chamaephytes are lumped together with hemicryptophytes as perennials (Fig. S2).

I used a dated molecular phylogeny of the British flora (Lim et al., 2014) available from TreeBASE (http://treebase.org/; accession number 15105). 14 species (3.5%) in the dataset were not present in the phylogeny. For 8 of these species, I used the position of a congeneric species as a proxy for the focal species (following Pennell et al., 2016). When multiple congeneric species were present, I consulted the phylogenetic literature to identify the most closely related proxy species (Scheen et al., 2004; Salmaki et al., 2013). For the remaining 6 missing species, I positioned them in the tree based on phylogenetic relationships to other genera or families present in the tree (Fior et al., 2006). Because many phylogenetic comparative methods do not allow polytomies, zero-length branches, and non-ultrametric trees, I made several small adjustments to the tree. I resolved polytomies randomly using the ‘multi2di’ function in **phytools** version 0.6-00 (Revell, 2012). I added 0.02 my to all zero-length branches, as this was approximately the length of the shortest nonzero branch length in the tree. After these changes, I slightly altered terminal branch lengths to make the tree precisely ultrametric.

I excluded C_4_ (3 species) and CAM (2 species) plants. I limited this investigation to angiosperms because only 4 non-angiosperms had stomata data. The final dataset contained 372 species (Fig. 1, S3). The R code accompanying this paper documents these decisions in greater detail and citations to the relevant literature.

**Figure 1:**
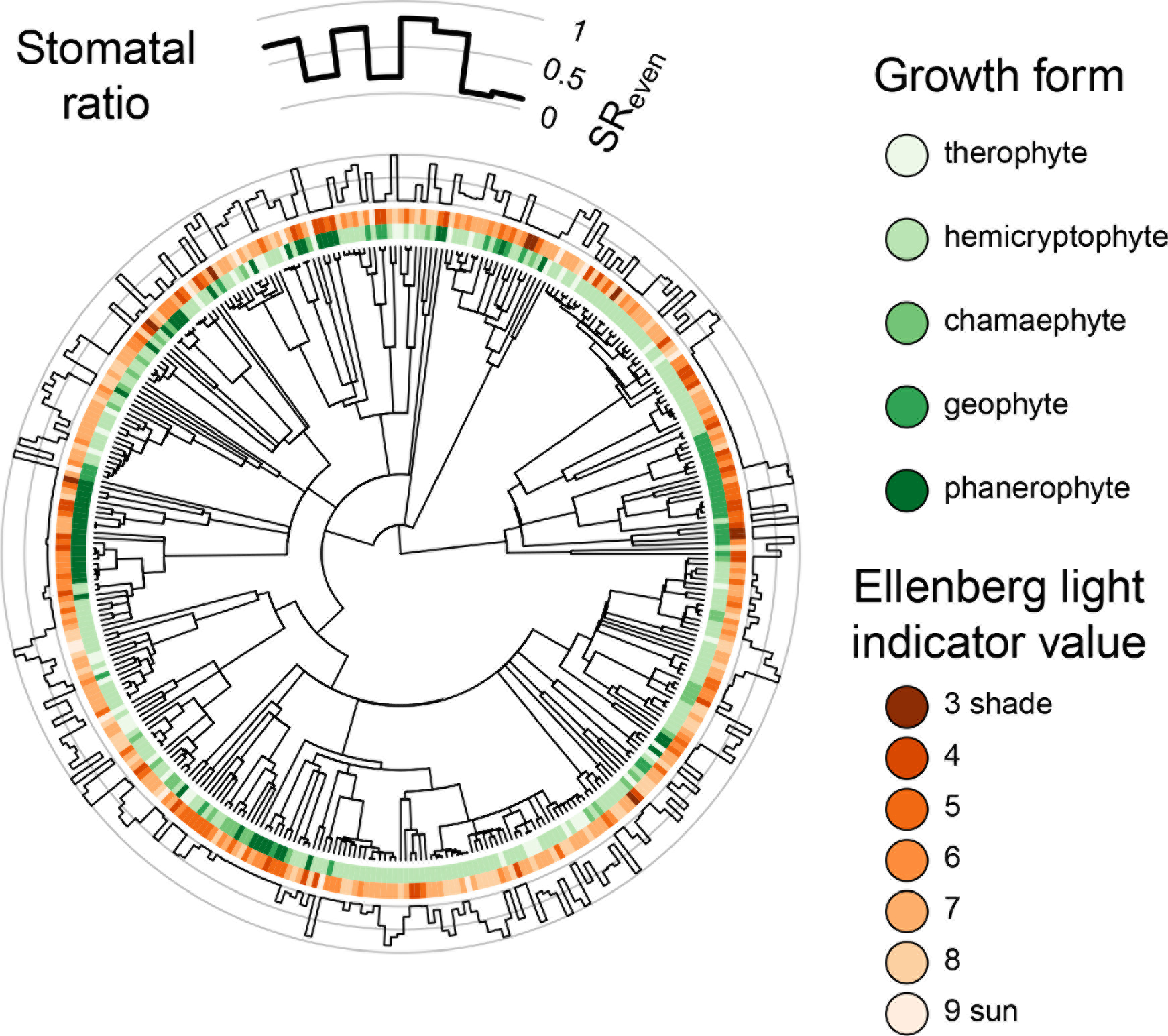
Phylogenetic diversification of stomatal ratio follows growth form and light tolerance. At the center is the phylogenetic tree for 372 species of British angiosperms. For each species, the green wedges indicate Raunkiær life form and the orange wedges indicate L-value. The outer circle indicates the stomatal ratio (SR_even_) for each species. As shown in the legend above, greater stomatal ratio means stomata are more evenly distributed across both leaf surfaces; lower stomatal ratio means that most stomata are on the lower surface.

Following Muir (2015), I calculated stomatal ratio in two different ways depending on what was most appropriate for the question:

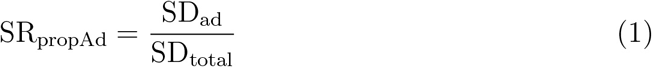

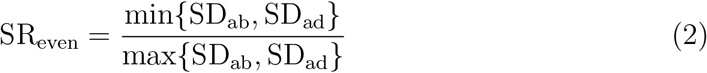

SD_ab_ and SD_ad_ are the stomatal densities on abaxial or adaxial surface, respectively. SD_total_ = SD_ab_ + SD_ad_. SR_propAd_ is the proportion of stomata density on the adaxial surface, which is useful for discriminating among hypostomatous (SR_propAd_ = 0), amphistomatous (0 < SR_propAd_ < 1), and hyperstomatous species (SR_propAd_ = 1). SR_even_ indicates how evenly stomatal densities are distributed across both leaf surfaces. This expression is useful because several hypotheses are based on the fact that a more even distribution should optimize leaf CO_2_ diffusion.

### Testing for an association between open habitat and growth form

I tested whether growth form, under either classification scheme, was associated with L-value among British angiosperms. I first used a phylogenetic ANOVA assuming an Ornstein-Uhlenbeck process model fit using **phylolm** version 2.5 (Ho and Ané, 2014). However, this analysis indicated no phylogenetic signal in the regression (See the R code accompanying this paper for further detail). Specifically, the estimated *α* parameter was extremely high. In the Ornstein-Uhlenbeck model, *α* is proportional to the inverse of the phylogenetic half-life (i.e. phylogenetic signal). When there is no phylogenetic signal (i.e. high *α*), regular and phylogenetic ANOVA converge on the same parameters estimates. Furthermore, statistical tests assuming there is phylogenetic signal when in fact none exists perform worse than nonphylogenetic tests (Revell, 2010). Therefore, I used a regular ANOVA with Type-2 sum of squares.

### Open habitat, growth form, and stomatal ratio

I compared phylogenetic linear models to test whether growth form, L-value, or interactions between them predicted SR_even_. I fit models using **phylolm** and calculated Akaike Information Criteria (AIC), a common measure of model fit that penalizes additional parameters. Phylogenetic linear models simultaneously estimate the effect of continuous and categorical predictors while controlling for phylogenetic nonindependence. For these and subsequent analyses, I assumed an Ornstein-Uhlenbeck process model for the residuals with the root character state integrated over the stationary distribution. The Ornstein-Uhlenbeck model is characterized by a diffusion rate (*σ*^2^) and a return rate (*α*), which describes the phylogenetic signal (see above). I used 10^4^ parametric bootstrap samples of the full model (including main effects and interactions) to calculate parameter confidence intervals (Boettiger et al., 2012).

I tested whether phylogenetic nonstationarity could explain the residual variation in stomatal ratio after accounting for growth form and L-value. Specifically, I compared the expected residual variation given the actual tree versus a hypothetical tree where trait evolution has reached stationarity (i.e. a star phylogeny with infinite branch lengths). If phylogeny explains much of the variation, then the simulated residual variance from the actual tree should be greater than that of the stationary tree. I simulated trait values from 10^4^ parametric bootstrap samples of the model with the lowest AIC (this was the model including Raunkiær life form, L-value, and their interaction; see Results). I performed the first set of simulations using the actual phylogenetic tree in **OUwie** version 1.50 (Beaulieu and O’Meara, 2016). Each simulation used a different bootstrap parameter sample of *α* and *σ*^2^, where *α* is the return rate to the mean and *σ*^2^ is the diffusion rate. At stationarity, the variance of an Ornstein-Uhlenbeck trait is equal to *σ*^2^/2*α*. Therefore, I simulated stationary data by assuming a normal distribution with this variance estimated from the bootstrap samples. For comparability, I set the mean of simulations from both actual phylogeny and the stationary ‘phylogeny’ to zero. I compared the actual to stationary variance across simulated datasets using a paired *t*-test.

### Does ab- or adaxial stomatal density contribute more to stomatal ratio evolution?

I used two related phylogenetic methods, variance decomposition and structural equation modeling (SEM), to assess the relative contribution of ab- versus adaxial stomatal density to light-mediated stomatal ratio evolution. First, the contribution of ab- versus adaxial stomatal density can be calculated using phylogenetic variance decomposition methods as derived below. Because stomatal density is highly skewed, I log-transformed values for normality:

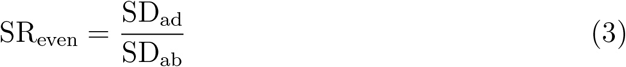

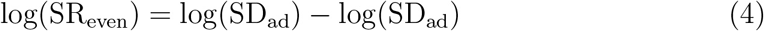

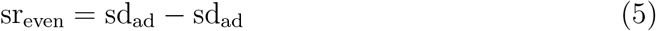

Lowercase variables (sr, sd) indicate log-transformed values. Because some species had zero adaxial stomata, I added one to all values prior to log-transformation. To make the variance decomposition calculations tractable, I have defined SR_even_ here as the ratio of ad- to abaxial stomatal density because in most cases adaxial stomatal density is lower than abaxial (see Eq. 2). This differs from analyses described above because in those I wanted to test what factors influenced the evenness of stomatal densities, regardless of which surface had higher density. With this modified form, the variance in sr_even_ can readily be decomposed into contributions of sd_ad_, sd_ab_, and their covariance:

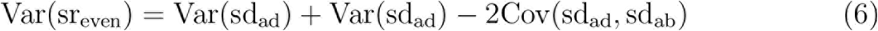

I did not use the raw covariance, but rather estimated the phylogenetic covariance matrix between L-value, sd_ab_, and sd_ad_ using a multivariate Ornstein-Uhlenbeck model fit in **Rphylopars** version 0.2.9 (Goolsby et al., 2016, 2017). The phylogenetic covariance measures how strongly a set of traits evolve together over macroevolutionary timescales. From the covariance matrix, I estimated the contribution of abaxial density, adaxial density, and their covariance as:

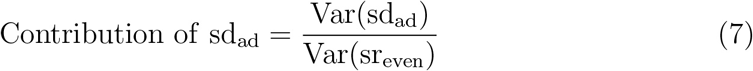

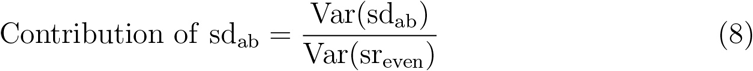

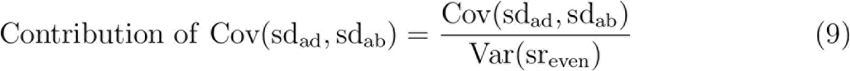

respectively. Note that when ab- and adaxial densities positively covary, the contribution will be negative because this reduces the variance in stomatal ratio.

If light-mediated increases in adaxial stomatal density can evolve while abaxial density remains roughly constant, then the phylogenetic regression of L-value on sd_ad_ will be stronger than that for sd_ab_. Under this scenario, stomatal ratio and density evolve in a coordinated fashion in response to light. Alternatively, if greater L-value favors greater stomatal ratio but total stomatal density is roughly constant, then there will be a negative covariance between ab- and adaxial density (Cov(sd_ad_, sd_ab_) < 0). I tested these competing predictions by fitting a simple phylogenetic SEM (see Mason et al., 2016, for a similar approach). In general, SEMs attempt to determine whether variables are related causally or whether a relationship is mediated by another correlated variable. Phylogenetic SEMs use the phylogenetic covariance matrix, as described above, rather than the raw covariance. Here, I used a phylogenetic SEM to simultaneously estimate regressions of L-value on sd_ad_ and sd_ab_ while allowing covariance between them (i.e. estimating Cov(sd_ad_, sd_ab_)). I used the R package **lavaan** version 0.5-23.1097 (Rosseel, 2012) to fit the SEM by finding parameter estimates would lead to phylogenetic covariance close to that observed in the data. I tested whether parameter estimates were significantly different from zero using *z*-scores.

## Results

### Light tolerance varies among growth forms

Ellenberg light indicator values (L-value) differed significantly among growth forms. Among Raunkiær life forms, therophytes (annuals), hemicryptophytes (perennial herbs with buds near the soil surface), and chamaephytes (subshrubs) had greater L-values than phanerophytes (woody plants) and geophytes (perennial herbs with storage organs) (Fig. 2; ANOVA - *F*_4,367_ = 18.3, *P* = 1.1 ×10^−13^). Likewise, herbaceous plants (annual, biennial, and perennials) had greater L-values than shrubs and trees (Fig. S4; ANOVA - *F*_4,367_ = 10.8, *P* = 2.6 × 10^−8^)

**Figure 2:**
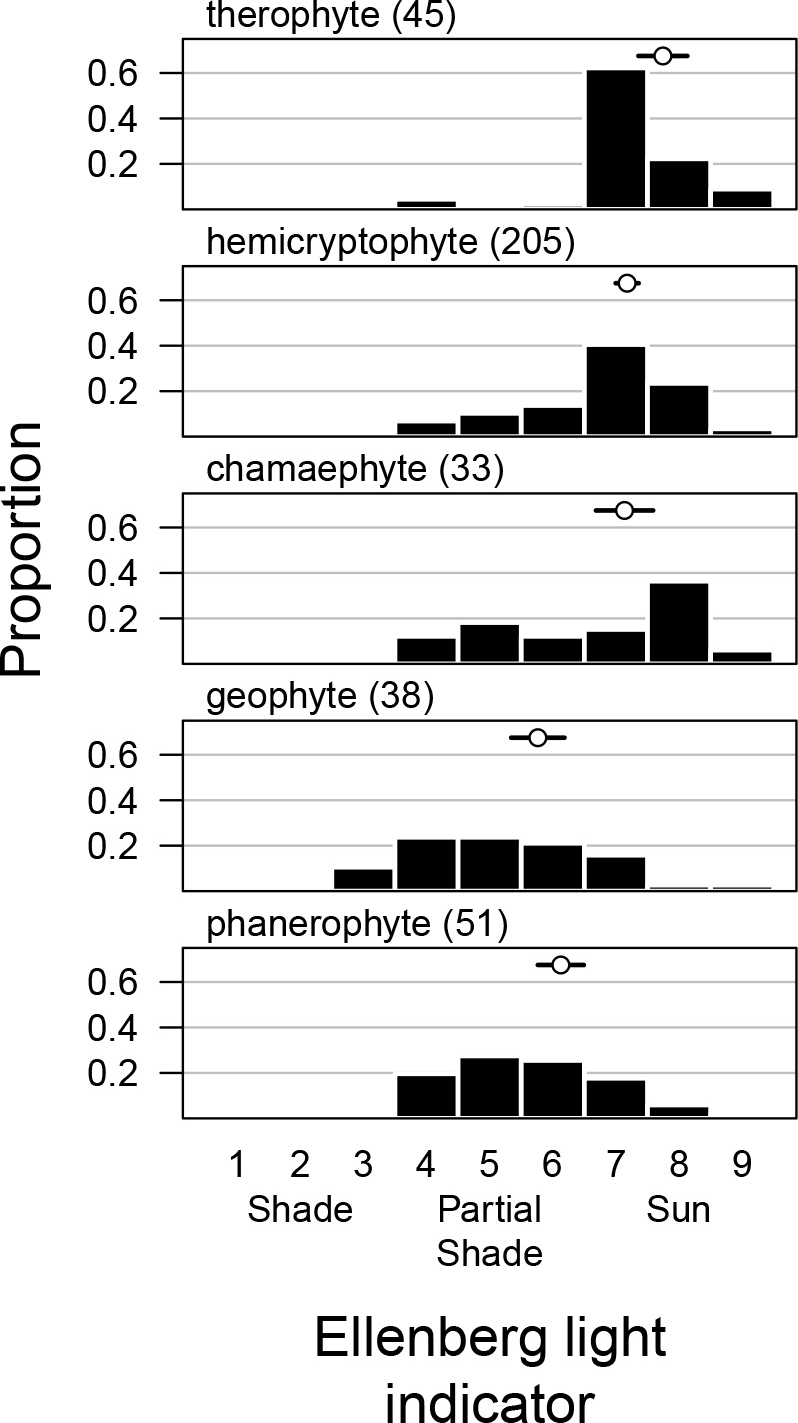
Life forms have different tolerances for sun and shade among British an-giosperms. Each panel is the distribution of Ellenberg light indicator values on an integer scale of 1-9 for different Raunkiær life forms. Height of the bars indicate the raw proportion of species in each bin for that life form. The sample size for each life form is listed next in parentheses. The mean (open circle) and 95% confidence intervals (black line) around the mean Ellenberg light indicator value for each life form based on phylogenetic regression are above the histogram.

### Interactions between light and growth form determine stomatal ratio

Overall, SR_even_ increased with L-value, but there was a significant interaction (ΔAIC > 2, Table 1) between Raunkiær life form and L-value (Fig. 3). When classified based on plant habit, growth form interacted with L-value less strongly (ΔAIC = 2.4; Fig. S5). Raunkiær life form explained variation in stomatal ratio better than habit (lower AIC; Table 1), therefore I focus hereafter on those analyses. Both life form and L-value significantly increased model fit, though L-value had a markedly larger effect on model AIC (Table 1). The significant interaction is caused by different slopes between life forms. Among life forms with the overall greatest L-value (therophytes, hemicryptophytes, and chamaephytes, see Fig. 2), there was a strong positive relationship between L-value and SR_even_. Parametrically bootstrapped 95% confidence intervals for the slope did not overlap zero (Fig. 3). The slope was weakly positive or not significantly different from zero in the most shade-adapted life forms (geophytes and phanerophytes), albeit the patterns were distinct in these groups. There were both hypostomatous (SR_even_ ≈ 0) and amphistomatous (SR_even_ ≈ 1) geophytes, but these were distributed across L-values. In contrast, phanerophytes were nearly always hypostomatous regardless of L-value.

**Table 1:** Interaction beween species’ Ellenberg light indicator value (L-value) and Raunkiær life form predict stomatal ratio (SR_even_). I compared phylogenetic linear models using the Akaike Information Criterion (AIC), where 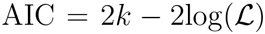. *k* is the number of model parameters and 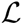 is the model likelihood. Given a set of candidate models, the difference in AIC between a model and the lowest AIC (ΔAIC) indicates the relative fit of competing models. The correlation coefficient *r*^2^ is another indicator of model fit. *α* and *σ*^2^ are the return rate and diffusion parameters of the Ornstein-Uhlenbeck model of trait evolution.

| Model: $SR_{\text{even}} \sim$ | $\alpha$ | $\sigma^2$ | $r^2$ | $k$ | $\log(\mathcal{L})$ | AIC | $\Delta AIC$ |
| --- | --- | --- | --- | --- | --- | --- | --- |
| L-value × Raunkiær life form | 0.46 | 0.068 | 0.34 | 12 | -33.3 | 90.6 | 0 |
| L-value × growth form | 0.46 | 0.07 | 0.32 | 12 | -38.2 | 100.3 | 9.8 |
| L-value + Raunkiær life form | 0.47 | 0.072 | 0.32 | 8 | -40.3 | 96.5 | 6 |
| L-value + growth form | 0.51 | 0.08 | 0.31 | 8 | -43.3 | 102.7 | 12.1 |
| Raunkiær life form | 0.34 | 0.067 | 0.15 | 7 | -79.2 | 172.4 | 81.8 |
| growth form | 0.35 | 0.069 | 0.13 | 7 | -82.5 | 178.9 | 88.4 |
| L-value | 0.64 | 0.108 | 0.26 | 4 | -59.3 | 126.6 | 36.1 |
| null | 0.29 | 0.067 | 0 | 3 | -107.5 | 221 | 130.5 |

**Figure 3:**
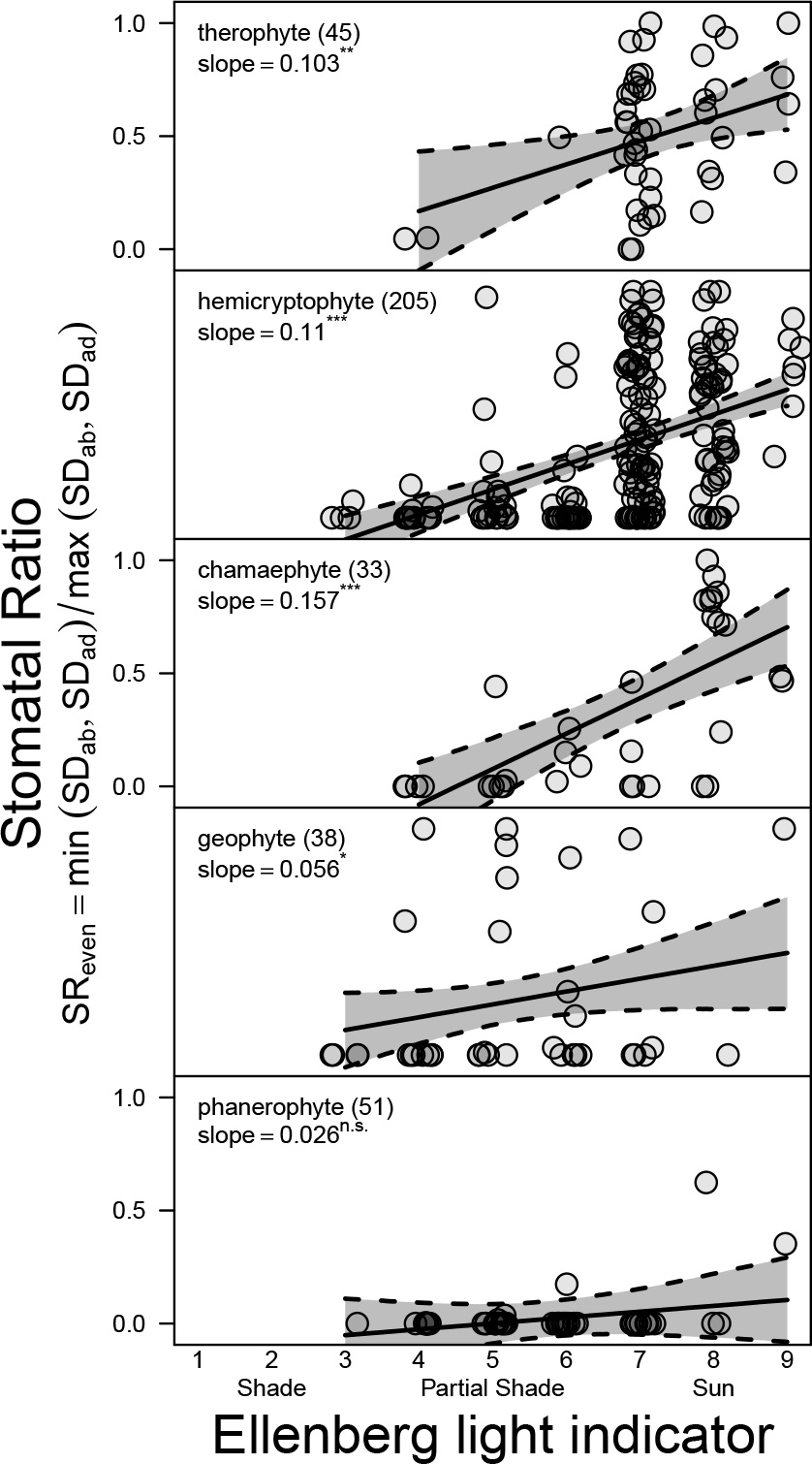
The effect of light on stomatal ratio depends on Raunkiær life form. Greater Ellenberg light indicator values (L-value) are associated with greater stomatal ratio (SR_even_) in therophytes, hemicryptophytes, and chamaephytes but not geophytes and phanerophytes. The maximum likelihood slope from phylogenetic regression is given with statistical significance based on 10^4^ parametric bootstrap samples. Numbers in parentheses next to Raunkiær life form are the sample sizes in the final dataset. Estimated slopes (solid line) and 95% bootstrapped confidence intervals (gray polygon between dashed lines) are plotted against raw data. Points have been jittered for visual clarity.

Although there was significant phylogenetic signal in the residual variation of stomatal ratio (see R code), the total variation among these species was consistent with a trait at stationarity. Specifically, the simulated residual trait variation, after accounting for Raunkiær life form and L-value, from the actual tree was not significantly greater than that simulated from a tree where traits had reached stationarity (paired *t*-test, *P* = 0.331). Hence, phylogenetic nonindependence is an important statistical consideration, but phylogeny does not explain stomatal trait variation among British angiosperms.

### Adaxial stomatal density contributes most of the variation in stomatal ratio

Adaxial (‘upper’) stomatal density contributed most to the evolutionary variation in stomatal ratio. The contributions of adaxial density, abaxial density, and their covariance are 1.14, 0.38, and −0.53, respectively. This implies that evolutionary variation in adaxial stomatal density is greater than that for stomatal ratio due to positive covariance between ab- and adaxial stomatal density.

Similarly, the phylogenetic SEM showed that changes in stomatal ratio associated with L-value can be attributed mostly to evolution of adaxial stomatal density (Fig. 4). Both sd_ad_ and sd_ab_ increased with L-value (*P* = 6.1 × 10^−7^ and 2.9 ×10^−5^, respectively). However, the regression of L-value on sd_ad_ was 2.1 × that of L-value on sd_ab_ (0.21 versus 0.1). Because stomatal densities were natural log-transformed, this implies an increase in L-value by one leads to a 1.23-fold change in adaxial stomatal density versus a 1.1-fold change in abaxial stomatal density. The SEM also showed a significant positive covariance between stomatal densities on each surface (*P* =1.7 ×10^−11^). These results together imply that total stomatal density increases with L-value, but the response is mediated mostly by increases in adaxial stomatal density.

**Figure 4:**
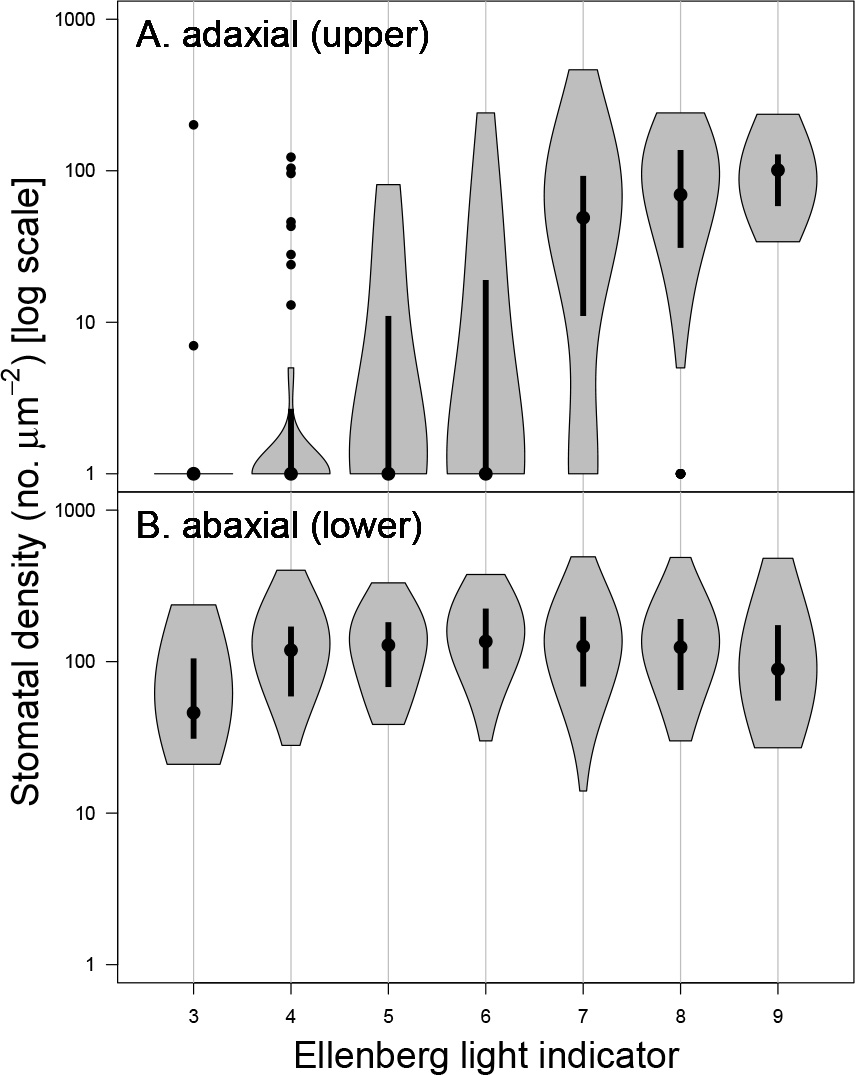
Light-mediated evolution of stomatal ratio is mostly driven by increased adaxial (‘upper’) stomatal density (Panel A), whereas abaxial (‘lower’) stomatal density (Panel B) is similar across Ellenberg light indicator values (L-value *x*-axis). The violin plot shows stomatal density (*y*-axis, log-scale) as a function of L-value. The width of the grey polygons indicates the density of data. Length of grey polygon indicate the range of the data; the dot indicates the median; the thick lines indicate the 0.25 and 0.75 quantiles. Points outside the polygons are statistical outliers.

## Discussion

The ratio of stomatal densities on the abaxial (‘lower’) to that of the adaxial (‘upper’) surface varies greatly across plant species, but the adaptive significance of this variation is not well understood. Comparative studies correlating stomatal ratio to ecological factors can distinguish among competing hypotheses and reveal critical experiments for future work. Previous comparative studies suggested that high light and herbaceous growth form favor amphistomy (Mott et al., 1984; Jordan et al., 2014; Muir, 2015; Bucher et al., 2017), particularly in the British flora (Salisbury, 1927; Peat and Fitter, 1994). However, none of these studies have accounted for the fact that light and growth form are often confounded – open, high light habitats are often dominated by herbs – or the fact that species are not independent because of shared evolutionary history. By bringing together datasets on stomata, light tolerance, growth form, and phylogeny of British angiosperms, I tested new hypotheses and reevaluated previous results using modern phylogenetic comparative methods. As expected, species’ light tolerance (Ellenberg light indicator or L-value) is confounded with growth form (Fig. 2, Fig. S4). Nevertheless, both L-value and growth form affect stomatal ratio, but these factors also interact. This new finding shows that the influence of L-value on stomatal ratio varies across forms. Finally, I show for the first time that adaxial stomatal density in particular accounts for most of the coordinated evolution between light tolerance and stomatal density. These novel findings provide further evidence that variation in stomatal ratio is adaptive and have important implications for interpreting changes in stomatal ratio through the paleo record (Jordan et al., 2014) and during domestication (Milla et al., 2013).

### Adaptive significance of amphistomy

Among British angiosperms, phylogenetic comparative analyses suggest that selection favors amphistomy in high light habitats among fast-growing herbs, but not shrubs and trees. This is a significant advance over previous studies that considered each factor in isolation and/or did not use modern approaches to control for phylogenetic nonindependence. For example, pioneering studies by Salisbury (1927) first suggested that amphistomy is associated with herbs in open habitats, albeit without formal statistical tools to disentangle light and growth form. Later work by Peat and Fitter (1994) demonstrated these trends again using family-level comparisons, a basic method to account for phylogenetic nonindependence (see also Mott et al., 1984; Beerling and Kelly, 1996). However, this approach is still problematic because traits like growth from can be highly phylogenetically conserved. For example, orders like Fagales contain multiple families dominated by hypostomatous trees, hence it is premature to conclude that this correlation is biologically meaningful without properly accounting for phylogenetic nonindependence. By combining trait, ecological, and phylogenetic datasets on British angiosperms, we now know that not only do both light and growth form influence stomatal ratio, but in fact their effects may reinforce one another. Based on information criteria, light may be a more important factor than growth form or their interaction (Table 1), consistent with previous studies indicating a dominant role of light (Mott et al., 1984; Jordan et al., 2014; Bucher et al., 2017).

The interaction between light and growth form among British angiosperms suggests that amphistomy may only be strongly favored when CO_2_ strongly limits photosynthesis (as in open habitat) *and* photosynthesis strongly limits fitness (as in herbs). This is consistent with life history theory predicting that the demography of open habitat herbs is strongly limited by plant growth (Franco and Silvertown, 1996). Along these lines, Raunkiær life form may explain stomatal ratio better than plant habit (Table 1) because it is a better proxy for life history characteristics. For example, on an axis of ‘fast’ to ‘slow’ life history, geophytes more closely resemble phanerophytes than do chamaephytes or hemicryptophytes (Salguero-Gómez et al., 2016). Similarly, the relationship between light and stomatal ratio for geophytes was intermediate between that for phanerophytes and chamaephytes/hemicryptophytes (Fig. S4). These comparisons indirectly suggest that both high light and fast life history are necessary to induce strong selection for amphistomy. The ideal way to test this would be to measure selection on stomatal ratio in a species that varied quantitatively in both stomatal ratio and life history (e.g., containing both therophyte/annual and perennial forms). I predict that amphistomy will be favored more strongly in the annual form grown under high light compared to an annual under low light or a perennial in high light, and much more strongly than a perennial grown in low light. Similar experiments could also be performed to test if and when light-mediated plasticity in stomatal ratio is adaptive (Gay and Hurd, 1975; Mott and Michaelson, 1991; Fontana et al., 2017).

The prevalence of amphistomatous species in high light habitats supports the hypothesis that amphistomy is an adaptation to maximize photosynthetic rates by increasing CO_2_ diffusion (Jones, 1985). It is also evidence against the hypothesis that the principle fitness cost of amphistomy is water loss (Darwin, 1886; Foster and Smith, 1986) or dehydration of pallisade mesophyll (Buckley et al., 2015), though these factors are likely very important in determining differential regulation of stomata on each surface. Since evaporative demand increases under high light, under these hypotheses we would expect plants in high light to be hypostomatous. Because stomatal conductances on each surface can be regulated independently in response to the environment (Darwin, 1898; Pospíŝilová and Solárová, 1984; Smith, 1981; Reich, 1984; Mott and O’Leary, 1984), amphistomatous leaves likely cope with these stresses by rapidly closing adaxial stomata when water supply cannot match evaporative demands (Richardson et al., 2017). Instead, patterns in the British flora are at least consistent with the idea that adaxial stomata increase susceptibility to foliar pathogens (Gutschick, 1984; McKown et al., 2014). The cost of adaxial stomata may be greater in the shade because wetter leaves and lower ultraviolet light provide a more suitable microclimate for many foliar pathogens.

### Amphistomy as a proxy for open habitat

These observations from the British flora partially support the hypothesis that amphistomy can be used a proxy for open habitat in paleoenvironment reconstruction (Carpenter, 1994; Jordan et al., 2014; Carpenter et al., 2015) but also point out previously unknown subtleties. These previous studies based their conclusions on data from Proteaceae, in which there is little quantitative variation in stomatal ratio; species are either completely hypostomatous (SR_propAd_ ≈ 0) or completely amphistomatous (SR_propAd_ ≈ 0.5). Stomatal ratio in British angiosperms is also bimodal (Peat and Fitter, 1994), but across many families there is also quantitative variation. Importantly, this means that quantitative variation in stomatal ratio may provide a more precise, quantitative indicator of vegetation type, rather than simply ‘open’ or ‘closed’. A quantitative relationship between L-value and stomatal ratio has already been shown for herbaceous perennials (Bucher et al., 2017), but we now know that it holds among annuals (therophytes), subshrubs (chamaephytes), and, to a lesser extent, geophytes as well (Fig. 3).

The weak or nonsignificant relationship between L-value and stomatal ratio in geophytes and phanerophytes suggests that in some cases amphistomy may not reliably indicate open habitat without further information. For example, perhaps amphistomatous geophytes from partially shaded habitats are spring ephemerals, so they experience high light during their growth phase, but this has not been tested. Likewise, phanerophytes are almost always hypostomatous (see also Muir, 2015). Most British phanerophytes are tall, hypostomatous trees, but the exceptions are telling. For example, the most amphistomatous phanerophyte in this dataset is *Brassica oleracea*, a short-statured biennial that has more in common physiologically with hemicryptophytes than other phanerophytes. The other amphistomatous phanerophytes in this data set (*Populus nigra* and *Lavatera arborea*) are fast-growing pioneer species.

Finally, phylogenetic information should improve inferences about paleoclimates because there is appreciable phylogenetic signal in stomatal ratio. The phylogenetic half-life of stomatal ratio evolution, after accounting for L-value and Raunkiær life form, is log(2)/*α* = 1.5 my (see Table 1 for maximum likelihood estimates of *α*, the return rate in the Ornstein-Uhlenbeck model of trait evolution). This lag time may indicate that evolving to the ‘optimum’ is constrained by the shape of the fitness landscape (Muir, 2015) or that other unmeasured factors which affect stomatal ratio have some phylogenetic signal. Regardless of the mechanism, this fact means that researchers may be able to use data from closely related species to improve paleoenvironment reconstruction. Despite there being phylogenetic signal, residual phylogenetic variation in stomatal ratio at the broad phylogenetic scale encompassed by British angiosperms should be at stationarity. The observed variance in stomatal ratio, after accounting for L-value and Raunkiær life form, was indistinguishable from that expected for a trait at stationarity under an Ornstein-Uhlenbeck process (see Results). This may not be the case for younger clades that have radiated in the past few million years.

### Coordinated evolution of stomatal ratio and density in response to light

Variation in stomatal ratio is determined primarily by evolution of adaxial stomatal density and is coordinated with increases in total leaf stomatal density summed across both surfaces. Note here that I am referring only to evolutionary variation in stomatal ratio among species; different processes may mediate within species variation or plastic responses. Phylogenetic analyses show that changes in stomatal ratio and total stomatal density, especially in response to L-value, are predominantly mediated by changes in adaxial stomatal density. To my knowledge, this highly nonrandom pattern among British angiosperms has not been demonstrated before, but it parallels evolutionary changes wrought by domestication (Milla et al., 2013); crop species tend to have higher adaxial stomatal density than their wild relatives.

There are at least two hypotheses that could explain why adaxial stomatal density is the most responsive. The first I refer to as the ‘real estate’ hypothesis. In hypostomatous plants, the lower surface is already crowded with stomata, and hence plants must increase the real estate available for stomata by developing them on the upper surface whenever there is selection for greater stomatal density. When stomata are packed too densely on one surface, stomatal interference limits their functioning and hence may create a strong selective pressure for amphistomy (Parlange and Waggoner, 1970; Dow et al., 2014).

I refer to the second hypothesis as the ‘coordination’ hypothesis. In this scenario, ecological conditions such as high light select for both increased total stomatal density and for amphistomy because these traits work well in coordination with one another. For example, if stomatal density were very high on a hypostomatous plant, then CO_2_ would be more strongly limited by the mesophyll. Adding a second parallel pathway for diffusion by developing stomata on both surfaces would restore a more optimal balance between stomatal and mesophyll limitations. Conversely, there would be little benefit to amphistomy when total stomatal density is low because CO_2_ diffusion is strongly limited by stomatal resistance, and therefore photosynthetic rate is not sensitive to changes in mesophyll diffusion mediated by stomatal ratio. A related prediction is that increased atmospheric CO_2_ may select for reduced stomatal ratio and density primarily by decreasing adaxial stomatal density, but this has not been well tested (but see Woodward and Bazzaz, 1988). These results suggest that coordination between stomatal ratio and density might play a greater role than previously appreciated in optimizing CO_2_ supply and demand under different light regimes (see also Beerling and Kelly, 1996).

### Conclusions

By revisiting this classic ecological dataset with modern phylogenetic comparative methods, I have shown that amphistomy is strongly associated with both light and growth form, but the interaction between these factors is also important. Furthermore, amphistomy and high stomatal density are closely connected in species from high light environments, suggesting selection for coordination between these traits.

## Acknowledgements

I thank Sally Otto, Matt Pennell, Rob Salguero-Gómez, and two anonymous reviewers for feedback on this manuscript. I was supported by an NSERC CREATE grant.

### Author contribution statement

CDM designed the study, collected data, analyzed the data, and wrote the manuscript.

## Supporting Information

**Figure S1:**
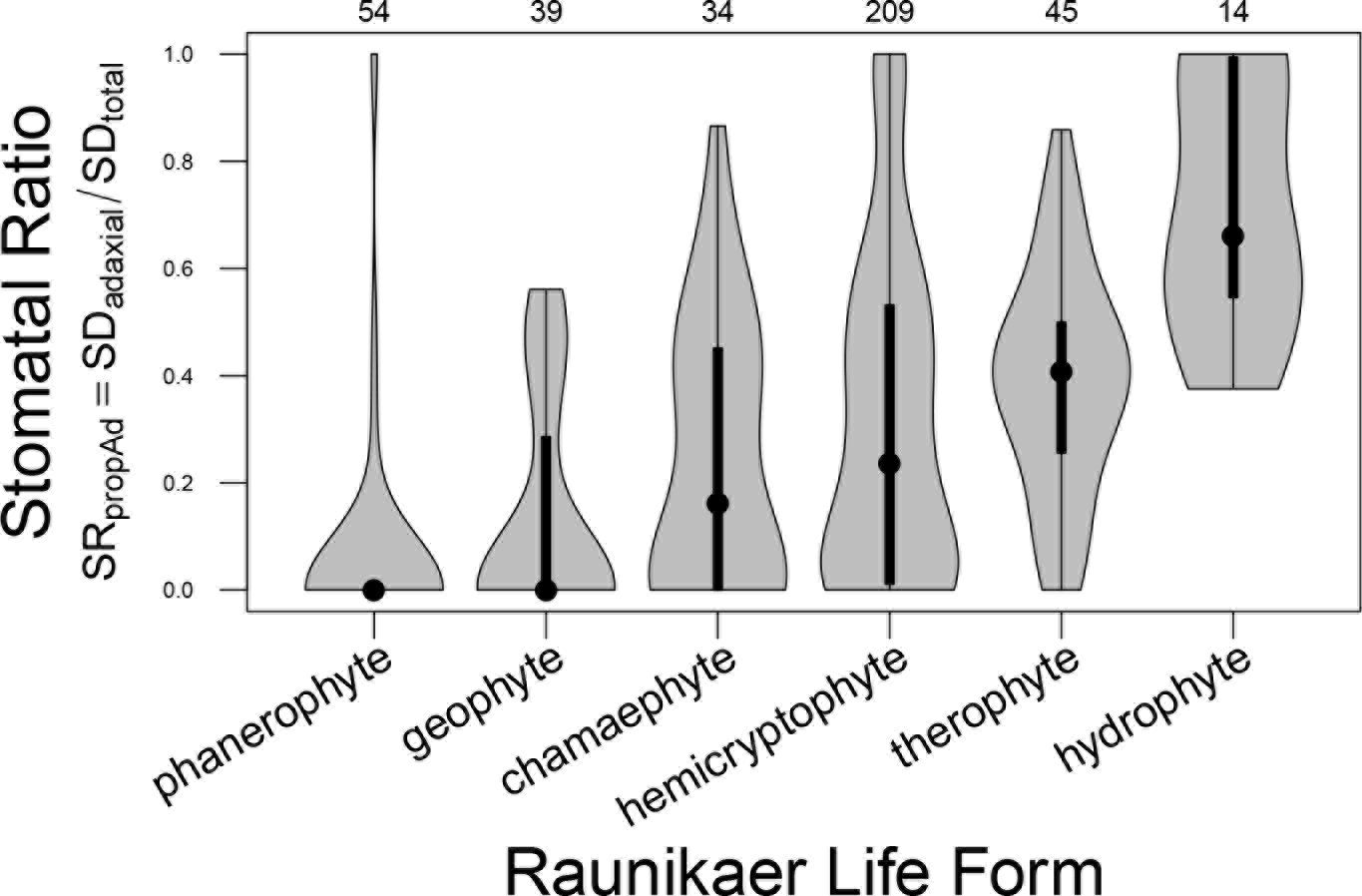
Most hydrophytes are hyperstomatous, having most stomata on the adaxial (‘upper’) surface (high SD_prop_Ad). The violin plot shows stomatal ratio as a function of Raunkiær life form. The width of the grey polygons indicates the density of data. Length of grey polygon indicate the range of the data; the point indicates the median; the thick lines indicate the 0.25 and 0.75 quantiles. Sample sizes per life form in the dataset are given above the upper plot margin. SD_ad_ and SD_total_ stand for the stomatal density on the adaxial surface and the total leaf surface (adaxial plus abaxial), respectively.

**Figure S2:**
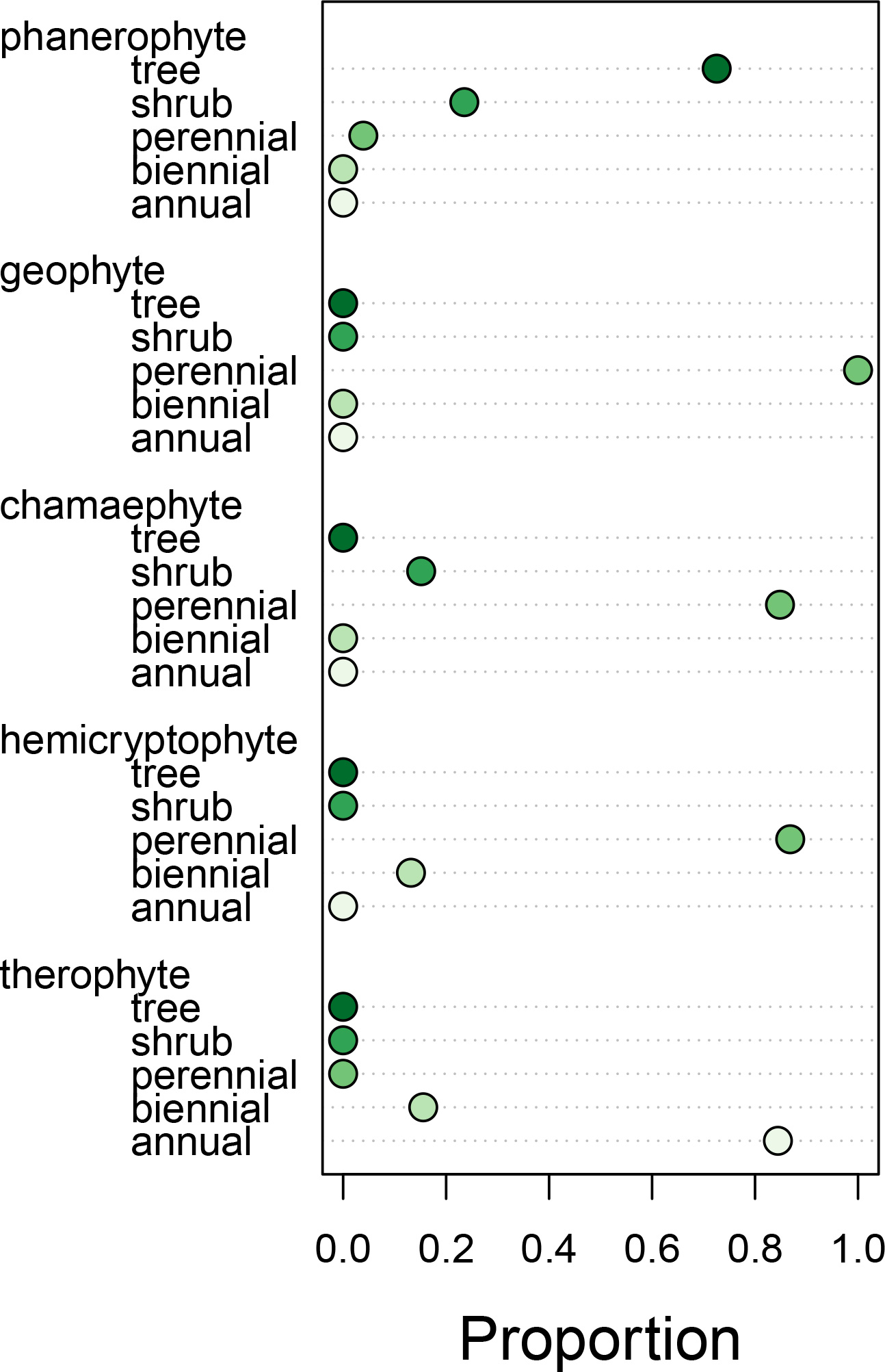
Raunkiær life form and plant habit broadly overlap. The dot chart shows for each Raunkiær life form, the proportion that overlap with a given plant habit. For example, phanerophytes are mostly trees and shrubs, geophytes are all perennial, therophytes are mostly annuals, and so forth.

**Figure S3:**
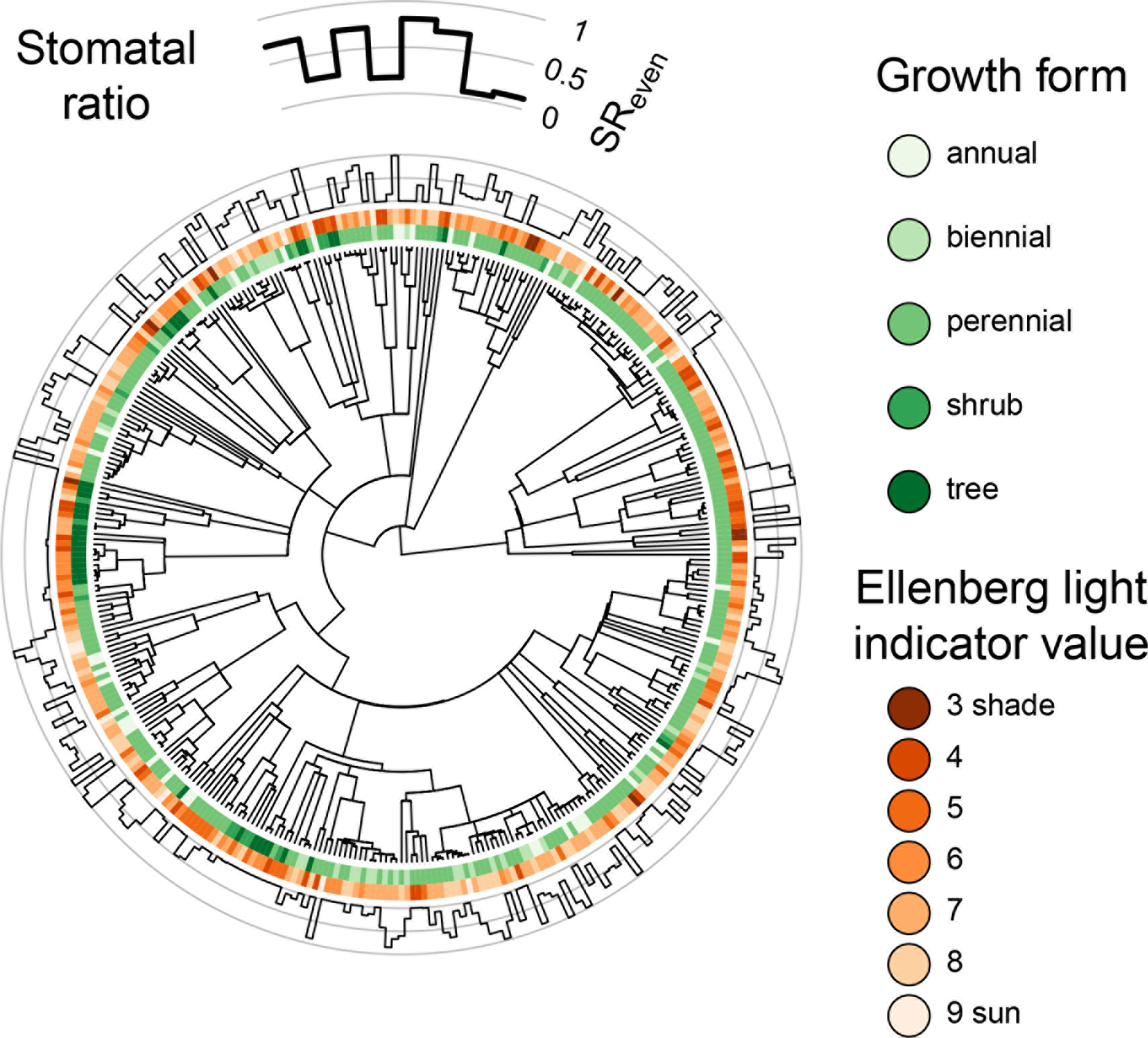
Phylogenetic diversification of stomatal ratio follows growth form and light tolerance. At the center is the phylogenetic tree for 372 species of British angiosperms. For each species, the green wedges indicate plant habit and the orange wedges indicate L-value. The outer circle indicates the stomatal ratio (SR_even_) for each species. As shown in the legend above, greater stomatal ratio means stomata are more evenly distributed across both leaf surfaces; lower stomatal ratio means that most stomata are on the lower surface.

**Figure S4:**
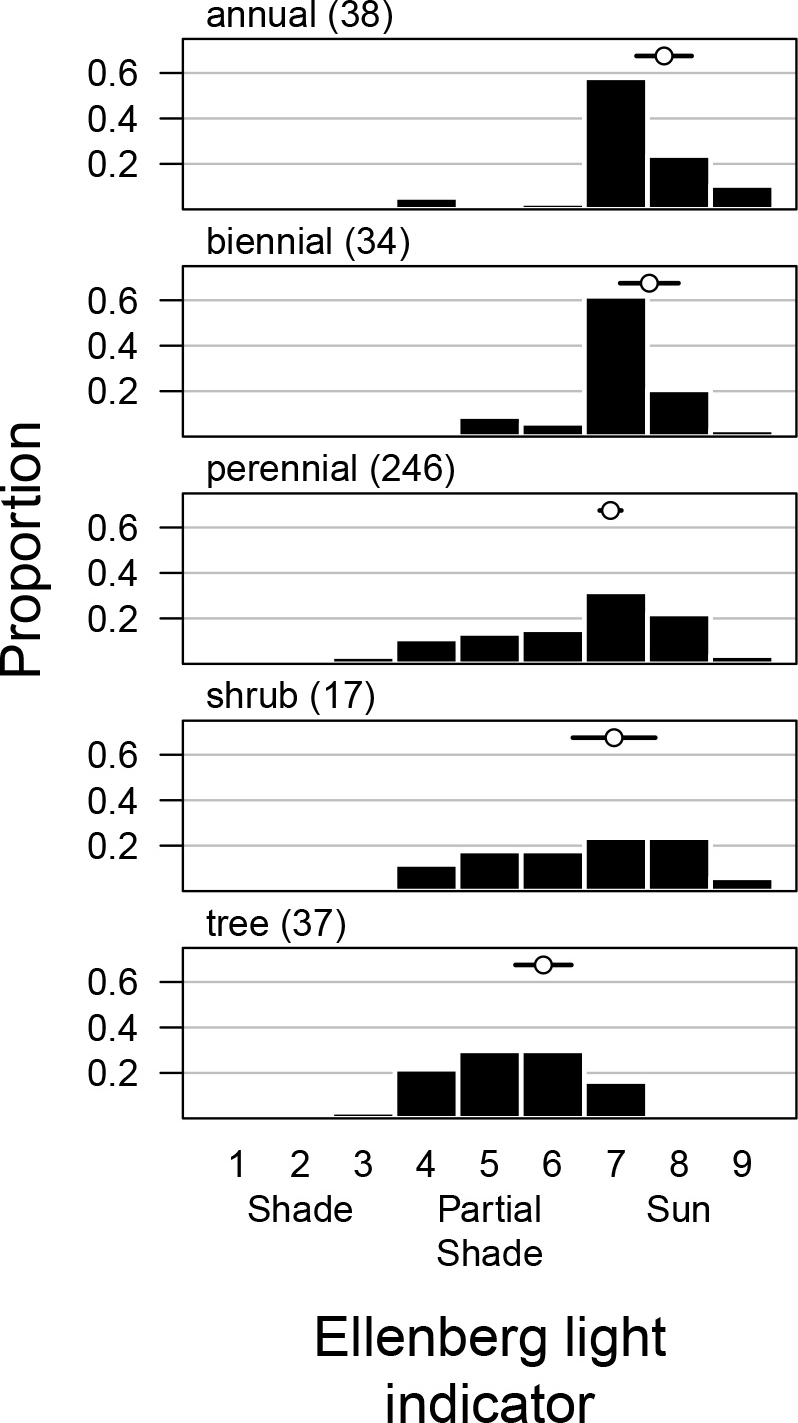
Growth forms have different tolerances for sun and shade among British angiosperms. Each panel is the distribution of Ellenberg light indicator values on an integer scale of 1-9 for different plant habits. Height of the bars indicate the raw proportion of species in each bin for that habit. The sample size for each habit is listed next in parentheses. The mean (open circle) and 95% confidence intervals (black line) around the mean Ellenberg light indicator value for each habit based on phylogenetic regression are above the histogram.

**Figure S5:**
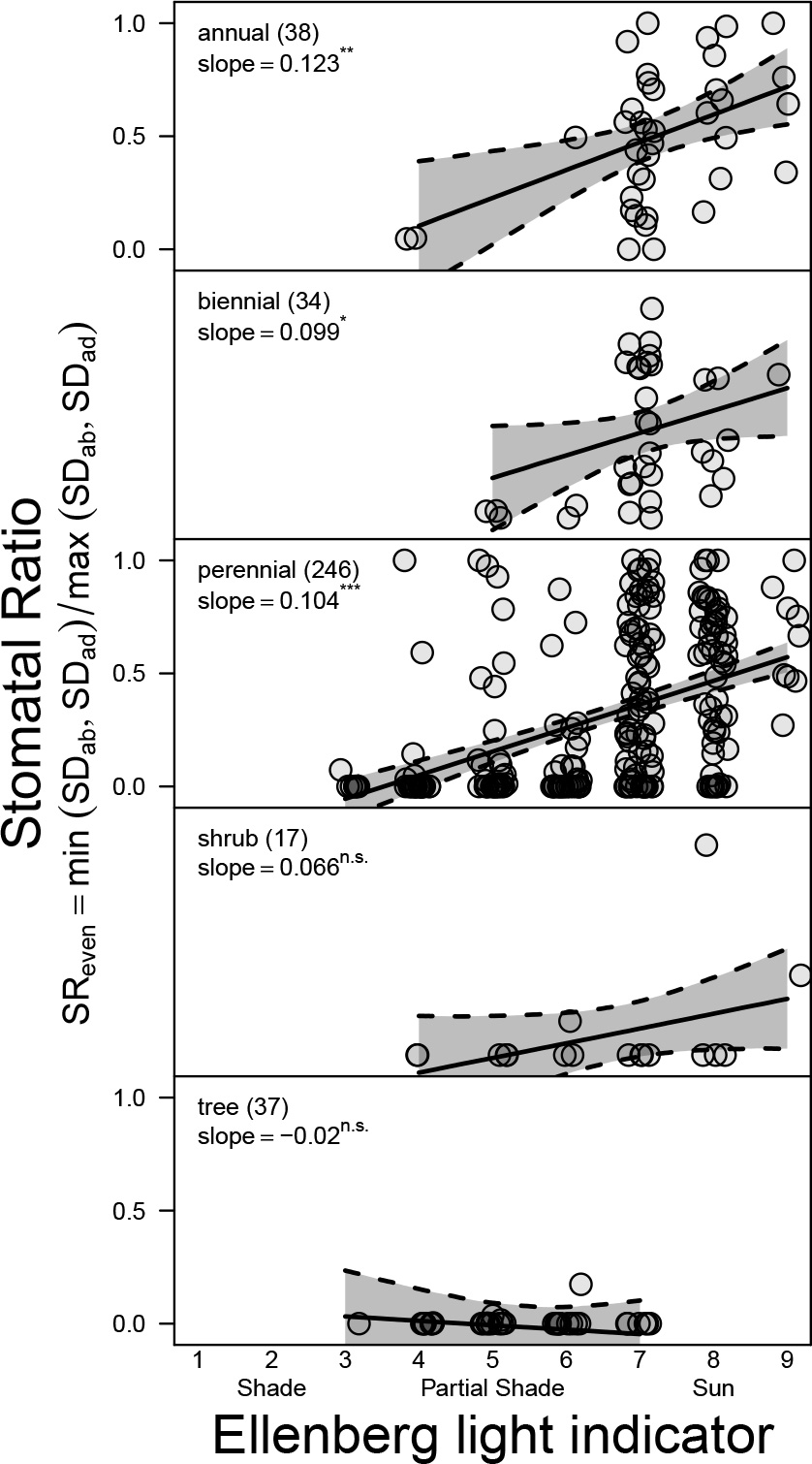
The effect of light on stomatal ratio depends on growth form. Greater Ellenberg light indicator values (L-value) are associated with greater stomatal ratio (SR_even_) in annual, biennual, and perennial herbs, but not shrubs or trees. The maximum likelihood slope from phylogenetic regression is given with statistical significance based on 10^4^ parametric bootstrap samples. Numbers in parentheses next to growth form are the sample sizes in the final dataset. Estimated slopes (solid line) and 95% bootstrapped confidence intervals (gray polygon between dashed lines) are plotted against raw data. Points have been jittered for visual clarity.

